# LRRK2 kinase dependent and independent function on endolysosomal repair promotes macrophage cell death

**DOI:** 10.1101/2023.09.27.559807

**Authors:** Rebecca Morrison, Simeon R Mihaylov, Chak Hon Luk, Angela Rodgers, Natalia Athanasiadi, Huw R Morris, Enrica Pellegrino, Sila Ultanir, Maximiliano G Gutierrez

**Affiliations:** Host-pathogen interactions in Tuberculosis Laboratory, The Francis Crick Institute, 1 Midland Road, London, NW1 1AT, United Kingdom; Kinases and Brain development Laboratory, The Francis Crick Institute, 1 Midland Road, London, NW1 1AT, United Kingdom; Department of Clinical and Movement Neurosciences, UCL Queen Square Institute of Neurology, UCL Movement Disorders Centre, University College London, London, UK

## Abstract

LRRK2 is commonly mutated in Parkinson’s disease and has cell type-specific mechanisms of activation and function. In macrophages, LRRK2 is associated with lysosomes and is activated following lysosomal damage. However, effects of pathogenic LRRK2-G2019S in macrophages are unknown. Here, using primary mouse and human iPSC-derived macrophage (iPSDM) models of LRRK2-G2019S, we defined the substrates of LRRK2 after lysosomal damage. Using phosphoproteomics we found that LRRK2-G2019S and wild-type macrophages showed similar levels of Rab phosphorylation after lysosomal damage, with the exceptions of Rab12 and Rab35, which were increased and decreased, respectively, in LRRK2-G2019S. LRRK2-G2019S macrophages showed a LRRK2 kinase activity-independent deficit in lysosomal membrane repair which resulted in more cell death and increased apoptosis. Importantly, we recapitulated this phenotype in iPSDM from patients carrying the G2019S mutation, but not in isogenic control iPSDM. Altogether, we define here the signaling downstream of G2019S in macrophages and identify susceptibility to cell death after lysosomal damage as an important phenotype of this mutation.

## Introduction

The leucine-rich repeat kinase 2 (LRRK2) gain-of-function G2019S mutation is the most common genetic cause of Parkinson’s disease (PD) (Ferreira and Massano, 2017; Paisan-Ruiz et al., 2004; Williams-Gray et al., 2006). The mutation is located in the kinase activation loop of LRRK2 and results in a modest 1.4 to 1.9-fold increase in kinase activity (Karayel et al., 2020; Myasnikov et al., 2021). LRRK2-G2019S is inherited in an autosomal dominant pattern with a relatively low age-related penetrance: at 79 years, 26% of carriers remain asymptomatic (Healy et al., 2008). To-date, it remains unclear what role LRRK2 has in cells and why overactive LRRK2-G2019S leads to disease. Key proteomic and phosphoproteomic studies in epithelial cells have revealed that LRRK2 kinase phosphorylates the switch-II motif of 14 Rab GTPases (Steger et al., 2017; Steger et al., 2016). While LRRK2 has been implicated in multiple cellular functions (Berwick et al., 2019), it is becoming increasingly clear that LRRK2 is important in lysosomal function (Bonet-Ponce and Cookson, 2022) and lysosomal membrane integrity and repair (Bonet-Ponce et al., 2020; Eguchi et al., 2018; Herbst et al., 2020).

LRRK2 is highly expressed by immune cells including macrophages (Gardet et al., 2010) and is expressed at higher levels in immune cells from PD patients (Cook et al., 2017). Innate immune cells such as macrophages can phagocytose extracellular substances including pathogens, cholesterol, urate crystals and in the context of PD, α-synuclein protein (Freeman et al., 2013). These substances may trigger lysosomal membrane damage, a process linked to inflammation and neurodegeneration (Lawrence and Zoncu, 2019; Papadopoulos and Meyer, 2017). Importantly, lysosomal membrane damage without efficient repair has been reported to result in different forms of cell death including apoptosis, necrosis and pyroptosis (Boya and Kroemer, 2008; Chen et al., 2019; Johansson et al., 2010; Radulovic et al., 2018; Skowyra et al., 2018). Endolysosomal damage triggers LRRK2 activation, Rab phosphorylation and endomembrane repair in murine macrophages, specifically RAW 264.7 cells (Herbst et al., 2020). However, the effects of LRRK2-G2019S mutation in this context remain unknown.

Here we sought to define the role of the kinase activity of LRRK2 and the most common pathogenic mutation G2019S in macrophage biology. Using genetic tools combined with proteomic and functional approaches we identified substrates that are phosphorylated by LRRK2 in macrophages. Importantly, we identified kinase activity-independent functions for LRRK2-mediated lysosomal membrane repair in macrophages. This deficiency in lysosomal repair led to a striking susceptibility to cell death. Our findings are relevant for the fundamental biology that links cell death and immune function of LRRK2 pathogenic mutations in macrophages.

## Results

### Identification of bona-fide LRRK2 substrates in macrophages

LRRK2 has a prominent role in macrophage function (Hartlova et al., 2018) and is activated after lysosomal damage (Herbst et al., 2020). However, the substrates that LRRK2 phosphorylates in macrophages after lysosomal damage are unknown. We generated bone marrow derived macrophages (BMDM) differentiated with GM-CSF from mice expressing LRRK2-G2019S (Tac-G2019S) and their respective wild-type (Tac-WT) (***see materials and methods***). In addition, as a control for LRRK2 kinase activity we generated BMDM from mice expressing a kinase-dead form of LRRK2 (NJ-D1994A) and their respective wild-type (NJ-WT). First, we confirmed macrophage differentiation by expression of surface markers and quantified lysosomal activity in basal conditions between the mutant BMDM (**fig. S1**). The surface expression of macrophage markers CD11b, F4/80, MHC II, TLR2, CD80, CD206 and CD11c was similar between the genotypes (**fig. S1, A and B**). Moreover, LysoTracker intensity levels, total LAMP-1 levels and proteolytic activity measured by DQ-BSA degradation showed no significant differences, indicating no lysosomal defect across the genotypes (**fig. S1, C to K**). Next, we utilised phosphoproteomics and the NJ-D1994A BMDMs to identify phosphorylation changes and the substrates of LRRK2 after lysosomal damage induced by the lysomotropic agent LLOMe. This approach revealed that a select group of Rab GTPases were phosphorylated by LRRK2 kinase after lysosomal damage: Rab3 pT86, Rab8 pT72, Rab10 pT73, Rab12 pS105, Rab35 pT72 and Rab43 pT80 (shown as red dots in **Fig. 1A to C**). Importantly, for these Rab GTPases, the total protein levels remained unaffected (**fig. S2, A and B**). While we did not detect a change in the phosphorylation of Rab44, its total level was increased in NJ-D1994A macrophages after LLOMe treatment (**fig. S2B**). Rab GTPase substrates containing the LRRK2 phosphorylation motif in their switch II domain that are known to be not phosphorylated by LRRK2 kinase (Rab1, Rab2 and Rab7a) (Steger et al., 2017; Steger et al., 2016) showed no significant changes between NJ-D1994A and NJ-WT BMDM after LLOMe-induced lysosomal damage (shown as green dots in **Fig. 1, A and B. fig. S3, A to C**). Additionally, the pS908 and pS910 sites on LRRK2 were less phosphorylated in the NJ-D1994A macrophages in both untreated and LLOMe-treated states, indicating that these phosphorylation sites were unaltered by lysosomal damage (shown as yellow dots **Fig. 1A and B, fig. S3 D and E**). Novel non-Rab proteins found to be altered in the NJ-D1994A BMDMs but unaffected by lysosomal damage included the proteins Mrpl51 and Mucl1 (phosphorylated less in NJ-D1994A macrophages) and Herpud2, Kctd12, Rpl21 and Rpl6 (phosphorylated more in NJ-D1994A macrophages) – suggesting basal levels of LRRK2 kinase activity (unaffected by activation status) may affect other non-Rab proteins either directly or indirectly (shown as blue dots in **Fig. 1A and B, fig. S3, F to M**). Alternatively, such consistent and LLOMe-independent phosphorylations could reflect genetic differences between the control and mutant mouse strains. Mutant mouse strain NJ-D1994A has the same genetic background with the control strain (NJ-WT) and has been backcrossed to the parent line two times with 98.1% homology on genetic monitoring (SNP analysis), yet it is possible that there are minor genetic differences. To further analyse specific changes in different conditions, we used Western blot to analyse the phosphorylation levels of the well-defined substrates under different conditions. Phosphorylation of Rab8, Rab10 and Rab12 protein was significantly higher after LLOMe-induced endolysosomal damage in NJ-WT mouse macrophages (**Fig. 1D to G**). We further confirmed this increase in pRab8, pRab10 and pRab12 was dependent on LRRK2 kinase activity as there was almost no phosphorylation in NJ-D1994A BMDM and we observed a significant reduction in phosphorylation levels in macrophages treated with MLi-2, a type I LRRK2 kinase inhibitor (**Fig. 1D to G**). Altogether, we found that in addition to Rab8, Rab10 and Rab12; the GTPases Rab3, Rab35 and Rab43 are substrates of LRRK2 in macrophages.

**Figure 1.**
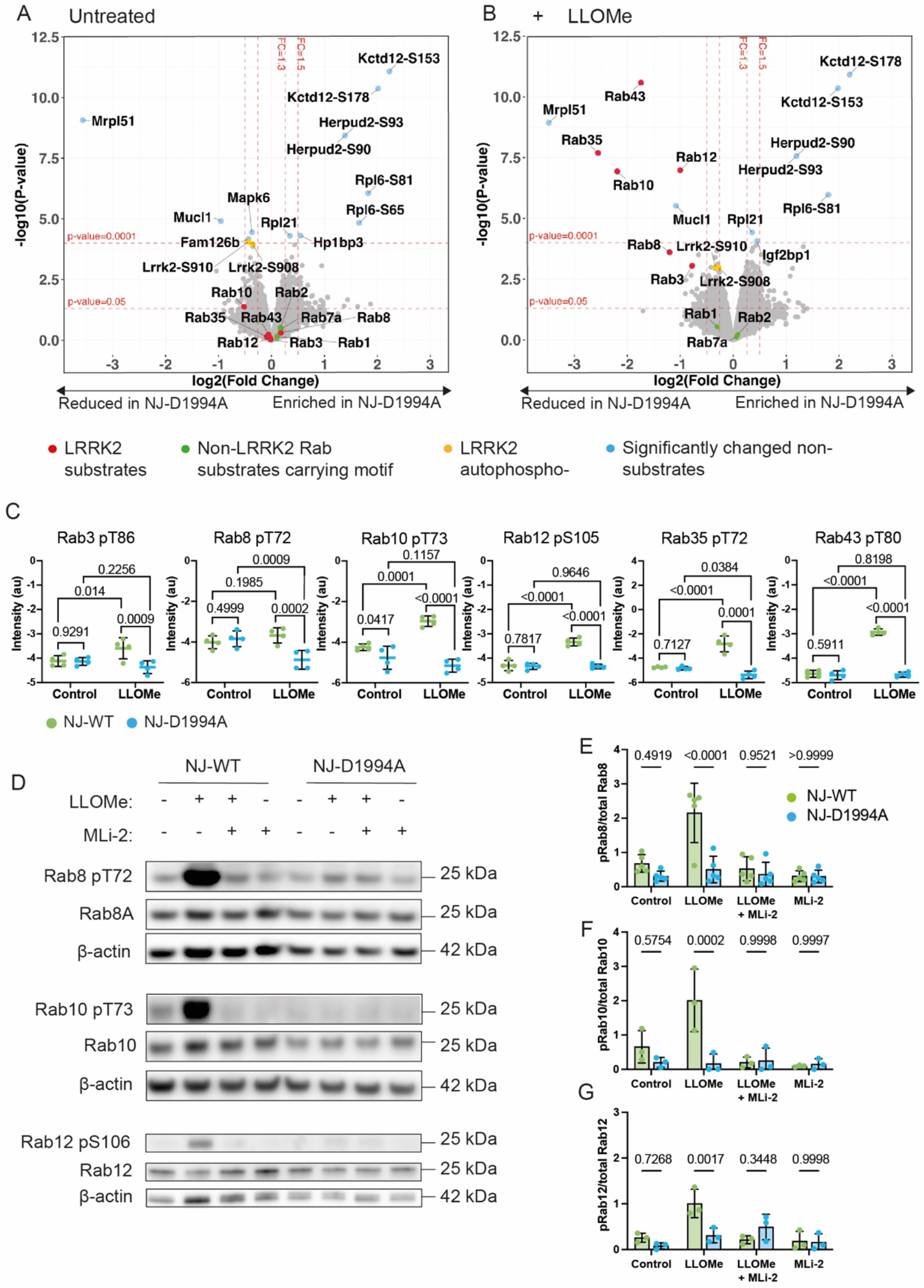
Identification of bona-fide LRRK2 substrates in macrophages. (A to B) BMDMs were treated with LLOMe 1 mM for 30 mins and cells were analysed by mass spectrometry. Volcano plots in the untreated (A) and LLOMe-treated (B) conditions of the difference in phosphorylation levels between NJ-WT and NJ-D1994A BMDMs. Each point represents one phosphorylation site of a protein. The x-axis shows the log_10_-transformed fold change, and the y-axis shows the significance by -log_10_-transformed P-value, obtained by linear models for microarray data. A fold-change greater than 1.3 and p value <0.05 was deemed significant. (C) Scatterplots showing the raw intensity data obtained by mass spectrometry and p-values for the significantly altered Rabs. Each point represents the normalised and log2 transformed TMT-corrected reporter intensity value obtained by mass spectrometry (n=4 biological replicates). P values by linear models for microarray data (D) Western blot analysis of Rab8 pT72, Rab8, Rab10 pT73, Rab10, Rab12 pS106, Rab12 and π-actin in NJ-WT and NJ-D1994A macrophages in untreated, MLi-2, LLOMe and LLOMe + MLi-2 conditions. (E to G) Rab8 pT72, Rab10 pT73 and Rab12 pS105 band intensities were quantified by densitometry and normalised to Rab8, Rab10 and Rab12, respectively. Data are mean ± SD from (E) 5 independent experiments and (F to G) 3 independent experiments. Two-way ANOVA followed by Šidák’s multiple comparisons test.

### Rab12 phosphorylation is upregulated by LRRK2-G2019S after lysosomal damage

We next tested whether overactive LRRK2-G2019S affects substrate phosphorylation after lysosomal damage using our phosphoproteomics approach. Unexpectedly, in the basal state, none of the Rab substrates showed significantly increased phosphorylation in Tac-G2019S compared to Tac-WT BMDMs (shown as red dots in **Fig. 2A**). Lysosomal damage induced by LLOMe resulted in increased phosphorylation of Rab12 pS105 in Tac-G2019S macrophages (**Fig. 2, A to C**). However, there was no difference in phosphorylation of Rab8 pT72, Rab10 pT73 or Rab43 pT80 between Tac-WT and Tac-G2019S macrophages and phosphorylation levels of Rab35 pT72 was in fact reduced in Tac-G2019S macrophages after LLOMe (shown as red dots in **Fig. 2B and C, fig. S3N**). In agreement with our results from the NJ-D1994A experiment, total levels of these Rab GTPases were unaltered at baseline and after LLOMe treatment, while total Rab44 levels were decreased after LLOMe in the Tac-G2019S macrophages (**fig. S2, C and D**). We identified altered phosphorylations in Tac-G2019S macrophages upon membrane damage and kinase activation: Cathepsin G and Hist1H3 both showed reduced phosphorylation in Tac-G2019S macrophages after membrane damage, while Mt2 and Dmxl2 showed increased phosphorylation in Tac-G2019S macrophages after membrane damage (shown as blue dots in **Figure 2B and C, fig. S3, O to R**). Cathepsin G enzyme is localized inside the lysosome and is not expected to be phosphorylated; its phosphorylation indicates a cytoplasmic localization due to lysosomal damage. Importantly, except for Mt2, these proteins did not show significant differences in total protein levels between Tac-WT and Tac-G2019S macrophages (**fig. S2, C and D**). These proteins were not identified as targets of LRRK2 kinase by our initial phosphoproteomics experiment (**Fig. 1, A and B**), suggesting possible indirect effects specific to the G2019S mutation only. Western blot analysis confirmed that levels of Rab12 pS105, but not Rab8 pT72 or Rab10 pT73, were increased in Tac-G2019S BMDMs after LLOMe compared to the wild-type strain (**Fig. 2D to G**). This effect could be reverted by treatment with MLi-2 (**Fig. 2, D and G**). Taken together, our findings suggest that in the G2019S genotype compared to the WT, there was not a striking effect in Rab GTPases phosphorylation except for Rab12, whose phosphorylation is increased by the G2019S mutation and only after lysosomal damage. It is interesting to note that LRRK2-G2019S also resulted in reduced phosphorylation of Rab35 after lysosomal damage, indicating that its pathogenic effects may not all be due to overactive kinase activity.

**Figure 2.**
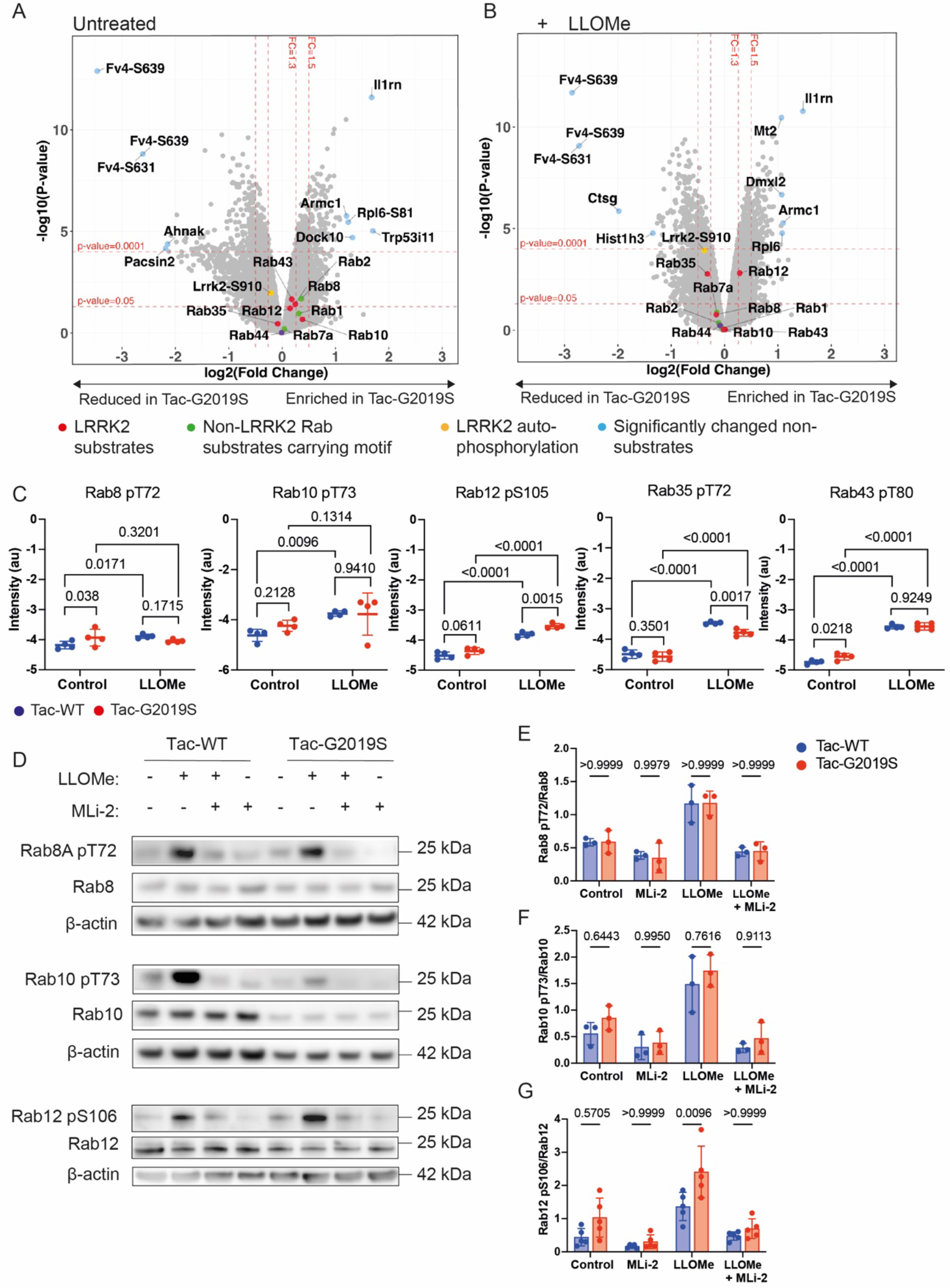
Rab12 phosphorylation is upregulated by LRRK2-G2019S after lysosomal damage in macrophages. (A to B) BMDMs were treated with LLOMe 1 mM for 30 minutes and cells were analysed by mass spectrometry. Volcano plots in the untreated (A) and LLOMe-treated (B) conditions of the difference in phosphorylation levels between Tac-WT and Tac-G2019S BMDMs. Each point represents one phosphorylation site of a protein. The x-axis shows the log_10_-transformed fold change, and the y-axis shows the significance by -log_10_-transformed P-value, obtained by linear models for microarray data. A fold-change greater than 1.3 and p value <0.05 was deemed significant. (C) Scatterplots showing the raw intensity data obtained by mass spectrometry and p-values for the Rabs which are phosphorylated by LRRK2 kinase as identified in the previous experiment. Each point represents the normalised and log2 transformed TMT-corrected reporter intensity value obtained by mass spectrometry (n=4 biological replicates) P values by linear models for microarray data (D) Western blot analysis of Rab8 pT72, Rab8, Rab10 pT73, Rab10, Rab12 pS106, Rab12 and β-actin in Tac-WT and Tac-G2019S macrophages in untreated, MLi-2, LLOMe and LLOMe + MLi-2 conditions. (E to G) Rab8 pT72, Rab10 pT73 and Rab12 pS105 band intensities were quantified by densitometry and normalised to Rab8, Rab10 and Rab12, respectively. Data are mean ± SD from (E to F) 3 independent experiments and (G) 5 independent experiments. Two-way ANOVA followed by Šidák’s multiple comparisons test.

### Repair of lysosomal damage is restricted in LRRK2-G2019S macrophages

In macrophages, LRRK2 is activated after lysosomal damage (Herbst et al., 2020) but the functional outcomes of lysosomal damage in a G2019S background are unknown. We aimed to study lysosomal damage and repair in LRRK2 mutant cells. To further understand this, we analysed lysosomal damage by measuring Galectin-3 (Gal3) positive lysosomes (Paz et al., 2010) in BMDM. We observed an increase in Gal3 positive vesicles after LLOMe treatment in both Tac-WT and Tac-G2019S macrophages (**fig. S4, A to C**). We did not observe significant differences in Tac-WT or Tac-G2019S macrophages treated with the kinase inhibitor MLi-2 (**fig. S4B**). Moreover, there were no differences between NJ-WT and NJ-D1994A macrophages (**fig. S4, A and C**), indicating that LRRK2-G2019S and LRRK2 kinase activity does not affect total levels of lysosomal damage. In contrast, when the component of the endosomal sorting complexes required for transport (ESCRT)-III complex CHMP4B localisation was analysed to monitor lysosomal repair, we found that the number of damaged lysosomes that recruited CHMP4B was significantly reduced in Tac-G2019S when compared to Tac-WT macrophages (**Fig. 3, A and B**). When cells were treated with MLi-2 the difference in CHMP4B recruitment between Tac-WT and Tac-G2019S macrophages was reverted, although overall levels of CHMP4B recruitment did not significantly change in Tac-WT or Tac-G2019S BMDM, suggesting that the recruitment is only partially dependent on the kinase activity of LRRK2 (**Fig. 3B**). This reduced CHMP4B recruitment was specific for LRRK2-G2019S since the kinase dead NJ-D1994A macrophages showed no differences in CHMP4B recruitment (**Fig. 3, A to C)**. These observations suggest that the effect in repair is specific to the G2019S mutation and partially dependent on the kinase activity of LRRK2. Confirming this result, the recruitment of ALIX, another ESCRT component, to damaged lysosomes was also reduced in Tac-G2019S macrophages (**Fig. 3, D and E**). To further confirm these observations, we used the lysomotropic dye LysoTracker to monitor lysosomal damage and repair by live cell snapshot imaging. While Tac-WT and Tac-G2019S macrophages leaked LysoTracker to a similar degree, the recovery of the lysosomal population after removal of LLOMe was reduced in Tac-G2019S macrophages (**Fig. 3, F and G**). Altogether, these findings indicate that endolysosomal repair is defective in LRRK2-G2019S macrophages.

**Figure 3.**
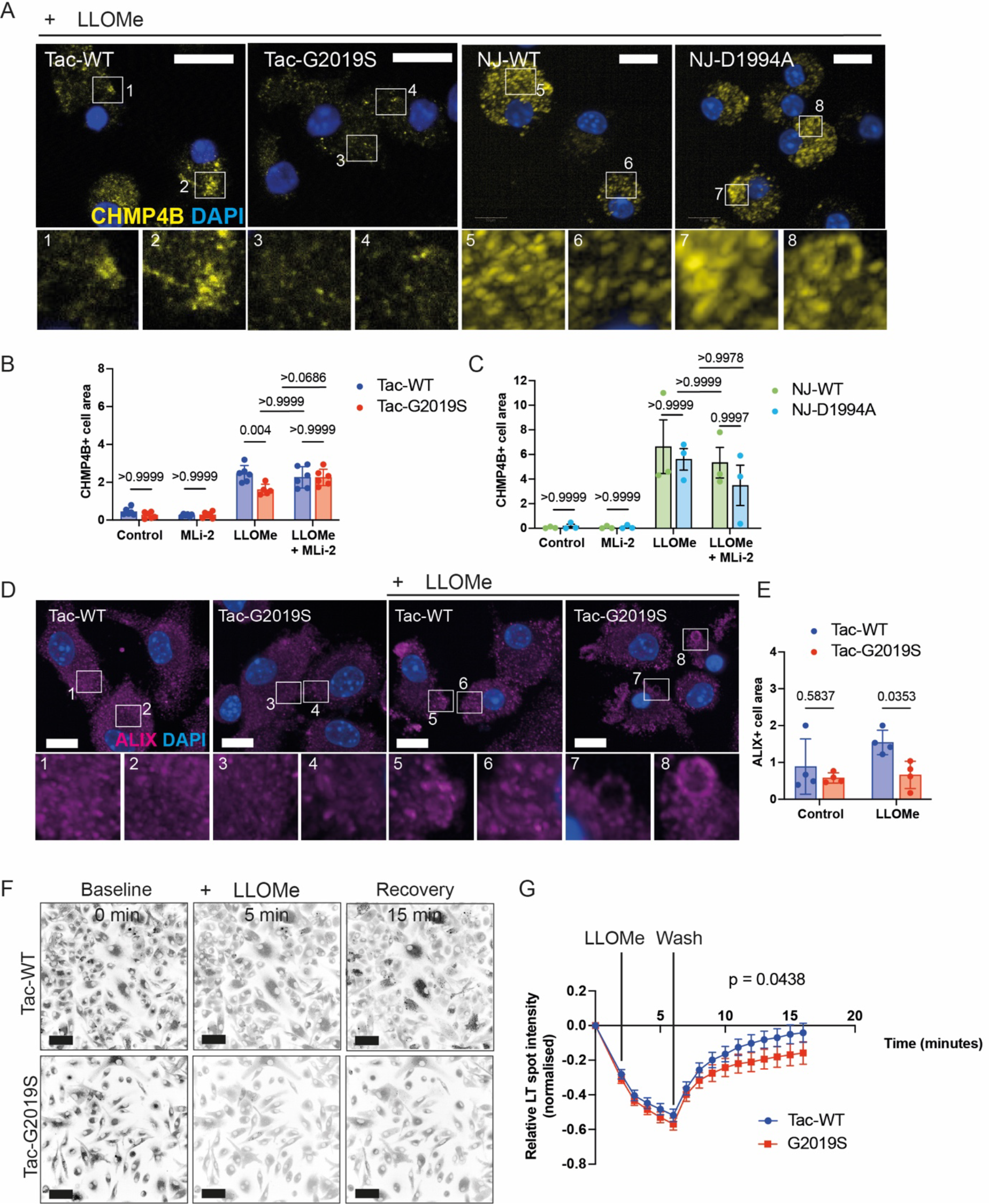
Endolysosomal damage repair is restricted in LRRK2-G2019S macrophages. (A to C) BMDMs were treated with LLOMe 1 mM for 30 minutes and endogenous CHMP4B positive cell area was visualised by immunofluorescence and quantified using Harmony software. Scale bars, 10 μm. Data representative of (B) two independent experiments (n=6 independent wells) and (C) 3 independent experiments (n=3 independent wells). Results are shown as mean ± SD. Two-way ANOVA followed by Šidák’s multiple comparisons test. (D to F) BMDMs were treated with LLOMe 1mM for 30 minutes and endogenous Alix positive cell area was visualised by immunofluorescence and quantified using Harmony software. Scale bars, 10 μm. Data representative of two independent experiments (n=4 independent wells). Results are shown as mean ± SD. Two-way ANOVA followed by Šidák’s multiple comparisons test. (G) Live cell snapshot imaging of LysoTracker positive spots in BMDM treated with LLOMe 1mM, followed by lysosomal recovery after removal of LLOMe and wash. Scale bar = 50 μm. (H) Data shown is the mean ± SD from 6 independent experiments. Differences between slopes in the recovery period (after wash) were estimated using linear regression.

### LRRK2-G2019S macrophages are more susceptible to cell death after lysosomal damage

Lysosomal damage without efficient repair will lead to cell death in epithelial cells (Radulovic et al., 2018), so we investigated if the observed deficiency in lysosomal membrane repair was associated with macrophage cell death. Strikingly, we found that Tac-G2019S macrophages showed significantly higher levels of cell death after LLOMe treatment (**Fig. 4, A and B**). In contrast, macrophages expressing the kinase dead variant of LRRK2 did not show differences in cell death (**Fig. 4, A and C)**. In line with this being a kinase activity-independent effect of LRRK2, the LRRK2-G2019S effect on cell death could not be reverted by treatment with MLi-2 (**Fig 4. D to F**). The increased susceptibility of Tac-G2019S macrophages to cell death was not due to differences in LLOMe processing as silica crystals, a membrane damage inducer independent of proteolytic activity, also induced more cell death in Tac-G2019S macrophages (**Fig. 4, G and H**).

**Figure 4.**
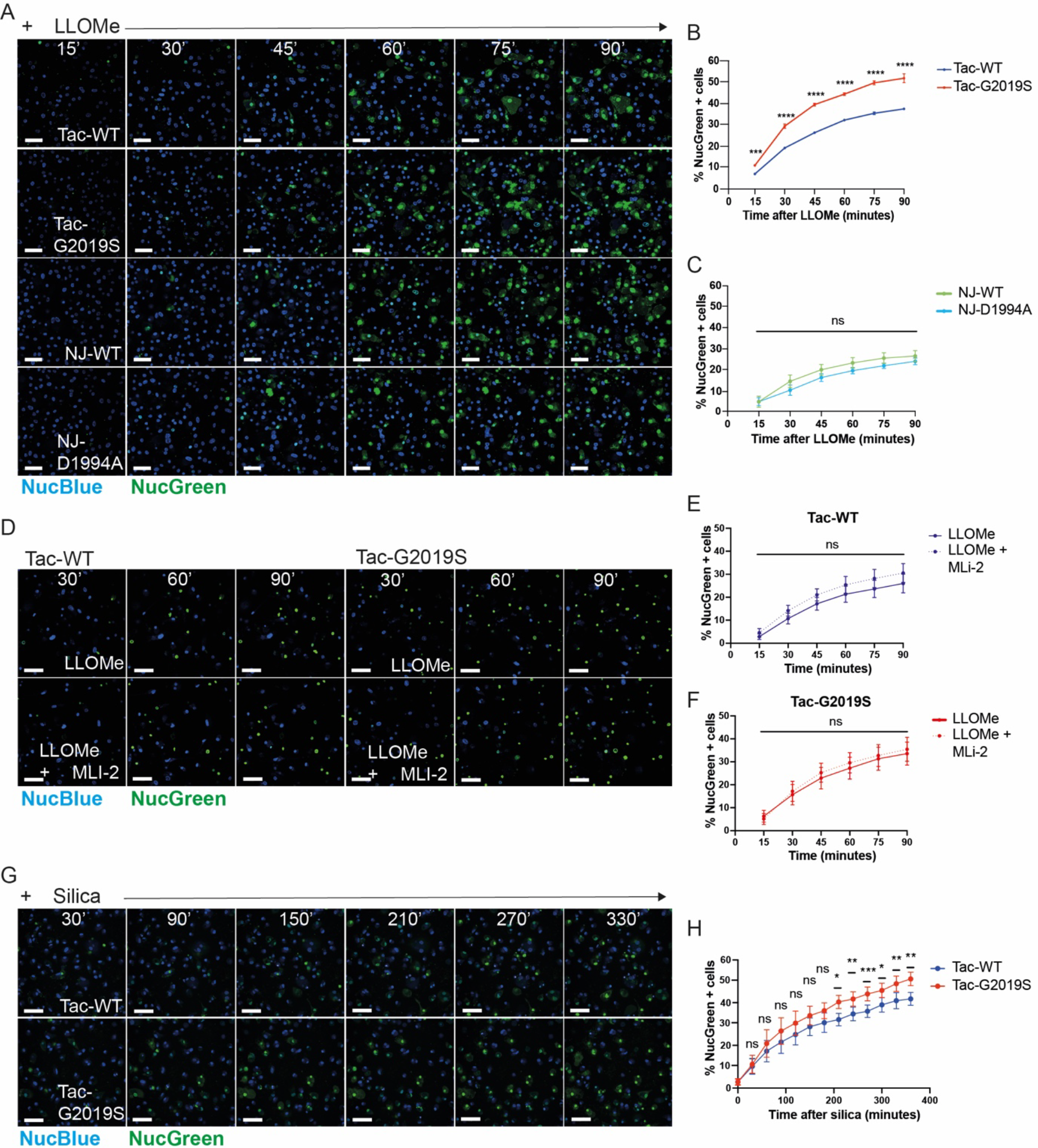
LRRK2-G2019S macrophages are more susceptible to cell death after lysosomal damage. (A) Snapshot of live BMDM treated with LLOMe 1 mM and imaged every 15 minutes. Nuclear staining for live/dead (blue/green). Scale bars = 50 μm. (B to C) Quantitative analysis of the percentage of dead cells at each timepoint. Two-way ANOVA followed by Šidák’s multiple comparisons test. (B) Data shown is the mean ± SD from one of four independent experiments (n=2 independent wells). (C) Data shown is the mean ± SEM from 3 independent experiments. (D) Snapshot of live BMDM pre-treated with MLi-2 100nM for 2h followed LLOMe 1mM and imaged every 15 minutes. Nuclear staining for live/dead (blue/green). Scale bars = 50 μm. (E to F) Quantitative analysis of the percentage of dead cells at each timepoint. Two-way ANOVA followed by Šidák’s multiple comparisons test. Data shown is the mean ± SD from 4 independent experiments (G) Snapshot of live BMDM treated with silica 300 μg/ml and imaged every 30 minutes. Nuclear staining for live/dead (blue/green). Scale bars = 50 μm. (H) Quantitative analysis of the percentage of dead cells at each timepoint. Data shown is the mean ± SD from 3 independent experiments (n=9 independent wells). Two-way ANOVA followed by Šidák’s multiple comparisons test. ns = non-significant; *P ≤ 0.05; **P ≤ 0.01; ***P ≤ 0.001; ****P ≤ 0.0001.

We then sought to explore the mode of cell death activated in Tac-G2019S and Tac-WT macrophages after LLOMe. Western blot analysis of markers of apoptosis (cleaved caspase-3 and cleaved PARP), pyroptosis (cleaved IL-1β and cleaved Gasdermin D) and necroptosis (phosphorylated RIP-3, phosphorylated MLKL, total RIP-3 and total MLKL) revealed that LLOMe treatment induced cleavage of PARP and caspase-3 and reduced total levels of RIP3 and MLKL in both Tac-WT and Tac-G2019S macrophages, indicating apoptosis and necrosis. When the inflammasome was primed by pre-treatment with LPS, LLOMe treatment resulted in cleavage of IL-1β and Gasdermin D in the macrophages, indicating that cell death by pyroptosis was also induced. As there was no phosphorylation of RIP-3 or MLKL detected, we concluded that necroptosis was not activated during lysosomal damage by LLOMe. There were no significant differences after 1 hour of LLOMe treatment in any marker of cell death (**Fig. 5, A to G**), but increased PARP and caspase-3 cleavage after 2 hours LLOMe in Tac-G2019S macrophages (**Fig. 5, B, I and K**). This suggests that the increase in cell death in Tac-G2019S macrophages is due to apoptosis. It has been reported that LPS-primed LRRK2-G2019S macrophages are more susceptible to reactive oxygen species (ROS) mediated necroptosis (Weindel et al., 2022). Therefore, we then primed macrophages with LPS and induced damage with LLOMe and monitored the levels of cell death by live cell snapshot imaging (**fig. S5)**. We found that while the levels of cell death increased with time of treatment, there were no differences between Tac-WT and Tac-G2019S macrophages, indicating that inflammasome activation does not contribute to the increased cell death seen in Tac-G2019S macrophages (**fig. S5**). Consistent with these data, there were no differences in markers of apoptosis, necroptosis or pyroptosis by Western blot in the LLOMe + LPS condition (**Fig. 5, A and B**). Cellular ROS are a known contributor to cell death pathways including apoptosis and necrosis (Redza-Dutordoir and Averill-Bates, 2016). Total ROS levels measured with CellROX were higher after lysosomal damage but similar between Tac-WT and Tac-G2019S macrophages (**fig. S6**). Altogether, these data argue that the G2019S mutation in LRRK2 results in a kinase activity-independent increase in cell death after LLOMe alongside an increase in cleaved PARP suggesting that this may be due to an increase in apoptosis.

**Figure 5.**
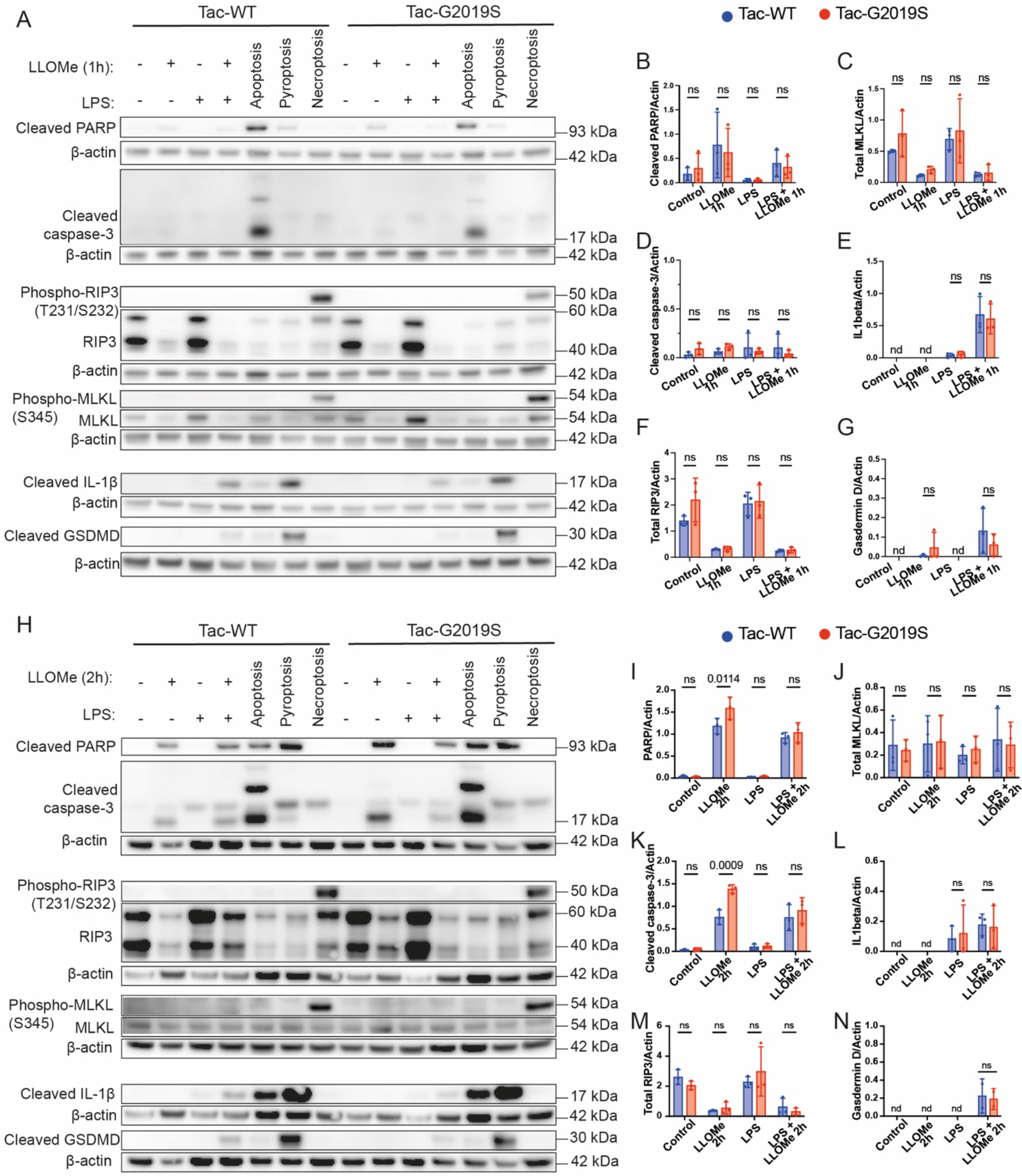
LRRK2-G2019S macrophages show increased PARP cleavage after 2h LLOMe. (A and H) BMDMs were pre-treated with LPS 250 ng/ml or vehicle for 3h followed by LLOMe 1 mM for the indicated time. For induction of apoptosis, BMDMs were treated with TNF-α 100 ng/ml and 5Z-7-Oxozeaenol 125 nM for 7h. For induction of necroptosis, BMDMs were treated with TNF-α 100 ng/ml, 5Z-7-Oxozeaenol 125 nM and Z-VAD-FMK 10 μM for 7 hours. For induction of pyroptosis, BMDMs were pre-treated with LPS 250 ng/ml for 3 hours followed by Nigericin 20 μM for 3 hours. Western blot for cleaved PARP, cleaved caspase-3, phospho-RIP3, RIP3, phospho-MLKL, MLKL, cleaved IL-1β, cleaved Gasdermin D (GSDMD) and β-actin levels. (B to G and H to N) Protein levels were quantified by densitometry and normalised to β-actin. Data represent the mean ± SD from 3 independent experiments. Two-way ANOVA followed by Šidák’s multiple comparisons test. nd = not-detected; ns = non-significant.

### iPSC-derived macrophages from LRRK2-G2019S patients are more susceptible to cell death after lysosomal damage

We next sought to validate our findings in human macrophages using a well-established iPSC-derived macrophage (iPSDM) model (Bernard et al., 2020; van Wilgenburg et al., 2013). iPSC derived from PD patients harbouring the G2019S mutation, obtained through the Michael J Fox Foundation PPMI resource, were gene-corrected to produce an isogenic control iPSC line using CRISPR/Cas9 genome editing (***see material and methods***). As a result, we obtained LRRK2-G2019S (PPMI ID clone: CDI00002173) and isogenic control (PPMI ID clone: 0043052321) iPSC. Both iPSC clones showed similar differentiation into iPSDM by expression of monocyte and macrophage markers and showed no differences in LysoTracker intensity or proteolytic activity measured by the DQ-BSA assay (**Fig. 6, A to E**). When we induced lysosomal damage with LLOMe we found significantly higher levels of cell death in G2019S iPSDM over time, confirming our observations in mouse macrophages (**Fig. 6, F and G**). This effect was independent of the kinase activity as MLi-2 did not revert the levels of cell death observed in the isogenic or G2019S iPSDM (**Fig. 6, F and H to J**). In line with the observations in BMDM, the effect on cell death observed in iPSDM could be attributed to a conformational change in LRRK2-G2019S that is not related to kinase activity. Altogether, we could recapitulate the findings in iPSDM after lysosomal damage and found that, like mouse macrophages, LRRK2-G2019S renders human macrophages more prone to cell death after lysosomal damage in a kinase activity-independent manner.

**Figure 6.**
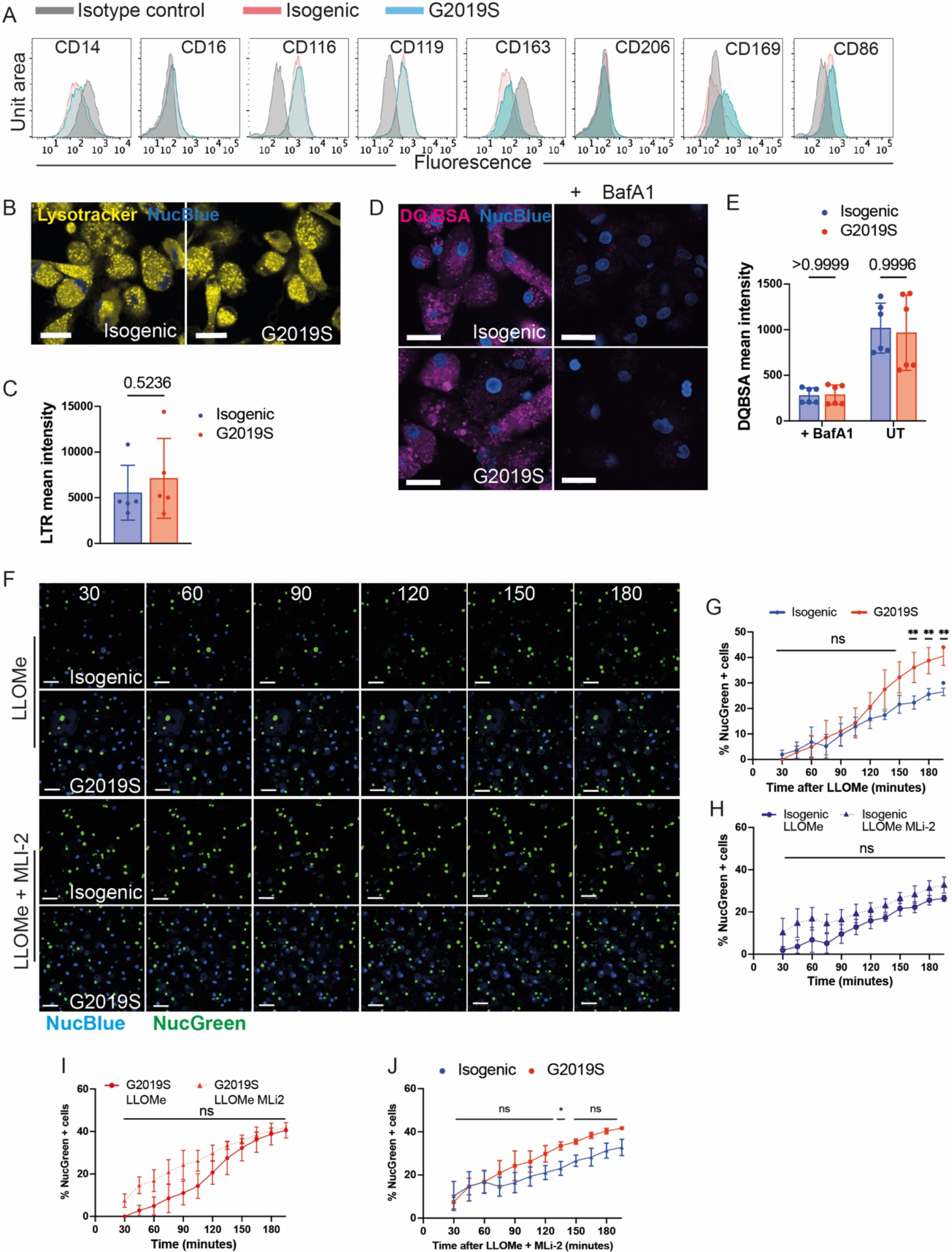
iPSC-derived macrophages from LRRK2-G2019S patients are more susceptible to cell death after lysosomal damage. (A) Flow cytometry characterisation of surface expression of macrophage and monocyte markers in iPSDM. Data is from one representative experiment. Data is presented as histograms with compensated fluorescence of the indicated surface marker on the x-axis. (B) Representative images of live iPSDM stained with Lysotracker (LTR) and NucBlue dye. Scale bars = 20 μm. (C) Quantitative analysis of the mean LTR cytoplasmic intensity. Data shown is mean ± SD from 2 independent experiments (n=5 independent wells). Two-tailed t-test. (D) Representative images of live iPSDM pre-treated with Bafilomycin A1 (BafA1) 100 nM followed by incubation with DQ-BSA and NucBlue dye. Scale bars = 20 μm. (E) Quantitative analysis of the mean DQ-BSA cytoplasmic intensity. Data shown is mean ± SD from 2 independent experiments (n=6 independent wells). Two-way ANOVA followed by Šidák’s multiple comparisons test. (F) Snapshot of live iPSDM treated with LLOMe 1 mM and imaged every 15 minutes. Cells were pre-treated with MLi-2 100 nM for 2 hours where indicated. Nuclear staining for live/dead (blue/green). Scale bars = 50 μm. (G to J) Quantitative analysis of the percentage of dead cells at each timepoint. Data shown is the mean ± SD from one representative experiment (n=3 independent wells). Two-way ANOVA followed by Šidák’s multiple comparisons test. ns = non-significant; *P≤0.05; **P ≤ 0.01.

## Discussion

The identification of Rab GTPases as the bona fide substrates of LRRK2 is ground-breaking progress in determining LRRK2 function in cells and in PD pathology (Steger et al., 2017; Steger et al., 2016). While studies have been expanded to many cell types, a holistic approach that identifies all substrates of endogenous LRRK2 in macrophages was lacking. It is likely that LRRK2 has cell type-specific functions, and this could be related to the subset of Rab GTPases expressed in different cell types. Since immune defects are thought to be critical for PD, and LRRK2 function is implicated in the immune response, we decided to address this gap. Here we detect a small group of Rab GTPases, Rab3, Rab8, Rab10, Rab12, Rab35 and Rab43, that are phosphorylated by LRRK2 after lysosomal damage in primary macrophages. Of these substrates, only Rab12 showed increased phosphorylation in G2019S macrophages, implicating Rab12 as an important mediator of LRRK2-G2019S in macrophages after membrane damage. These results agree with studies that identified Rab12 as a key activator of LRRK2 kinase activity in epithelial cells (Herschel et al., 2023; Vitaliy et al., 2023). The function of Rab12 in macrophages is unknown. However, Rab12 has been linked to phagocytosis (Gutierrez, 2013) and inflammation (Prashar et al., 2017). Defining the function of Rab12 in macrophages and the link to LRRK2 activity require further studies. While total levels of most Rab GTPases were unaltered by LLOMe treatment or LRRK2 mutations, Rab44 levels robustly increased after LLOMe in D1994A macrophages and significantly decreased after LLOMe in G2019S macrophages. Rab44 is an atypical Rab GTPase which is much larger than other Rab GTPases (molecular weight 110 kDa) containing additional domains (Yamaguchi et al., 2018). Intriguingly, Rab44 has been linked to lysosomal exocytosis, a process that may be triggered by lysosomal damage, in mast cells (Kadowaki et al., 2021). Further studies into the role of Rab44 in lysosomal damage and how LRRK2 mutations could affect expression are required. Rab proteins that have not been detected in our experiments, could be expressed and modulated by LRRK2 and/or membrane damage. Mass spectrometry may not detect all expressed proteins, therefore we cannot reach conclusions about the Rabs that have not been detected.

Given previous data indicating a 1.4 to 1.9-fold increase in LRRK2-G2019S kinase activity as compared to wild type (Karayel et al., 2020; Myasnikov et al., 2021), it was unexpected to find that Rab GTPase phosphorylation was not significantly increased in basal conditions in G2019S macrophages. In contrast, our findings indicate that phosphorylation of Rab35 is slightly reduced in G2019S macrophages after lysosomal membrane damage, while phosphorylation levels of Rab8, Rab10 and Rab43 were similar between WT and G2019S macrophages. Further, there was reduced phosphorylation of Cathepsin G and the histone Hist1H3 after membrane damage in G2019S macrophages – the downstream consequences of this may be of interest for future studies. Importantly, our experiments were performed in cells under endogenous LRRK2 expression and not by using over expression. Our findings highlight that LRRK2-G2019S kinase activity does not translate to a pronounced phosphorylation effect in primary macrophages; and we postulate that LRRK2-G2019S may not result in a widespread overactivity in cells.

LRRK2 is implicated in lysosomal function, and it is activated after lysosomal damage. There are reports in different cell types showing that proteolytic activity and lysosomal function is impacted by the G2019S mutation (Henry et al., 2015; Narayana and Shawn, 2023). Unexpectedly, we found here that both LRRK2 kinase dead and G2019S primary mouse bone marrow-derived macrophages do not show a significant change in proteolytic activity by using at least two functional assays. In agreement, we do not see major changes in lysosomal enzyme content by proteomics across the different genotypes. It is likely that the differentiation approaches to generate cells *in vitro* could affect this as it is known that myelocytic cells have qualitative and quantitative differences in lysosomal content (Delamarre et al., 2005).

LRRK2 is activated after lysosomal damage, and it is implicated in lysosomal repair by the ESCRT-III mediated machinery and likely other vesicle repair mechanisms. We show here that macrophages expressing the pathogenic mutation G2019S are not able to efficiently repair lysosomes after damage with reduced ESCRT recruitment. Surprisingly, this repair defect was only partially kinase activity independent as MLi-2 was only partially able to revert the effect and the kinase dead mutant had no effect. There is compelling evidence that mutations in LRRK2 could affect conformational changes and lead to interaction with other proteins or cytoskeleton components (Taylor and Alessi, 2020). This is the reason why there are current developments of type II inhibitors that target open/close conformational changes in LRRK2 (Tasegian et al., 2021) and PROTACs that target LRRK2 to degradation (Konstantinidou et al., 2021). The G2019S mutation is not very penetrant, and phenotypes *in vitro* are predicted to be mild. Our data suggest that conformational changes in LRRK2 are the result of the mutation, as these changes will affect downstream factors regulating membrane repair. Our studies in macrophages therefore highlight kinase independent functional effects of LRRK2 in macrophages.

Overexpression of mutant LRRK2-G2019S in cell culture systems is associated with neurotoxicity (West et al., 2007), whereas overexpression of kinase-dead LRRK2 is protective against neurotoxicity (Greggio et al., 2006). LRRK2-G2019S has been reported to link mitochondrial homeostasis and necroptosis mediated by Gasdermin D (Weindel et al., 2020; Weindel et al., 2022). In our experiments, we did not observe cell death alterations after LPS priming or differences in markers of necroptosis or total levels of ROS. It is possible that different mouse strains used in these studies as well as differentiation protocols (e.g., different growth factors) account for the differences observed here. In fact, we observed differences in basal levels of phosphorylation of Rab GTPases in the different WT strains (data not shown), indicating that direct comparisons between different genetic backgrounds will be difficult. Further, our data indicate that there is no activation of necroptosis following lysosomal damage, which would explain why we do not recapitulate this phenotype here. Our results show increased poly-ADP ribose polymerase (PARP) cleavage in G2019S macrophages after membrane damage. PARP cleavage is implicated in multiple forms of cell death including apoptosis, necrosis and a recently described form of cell death known as parthanatos (Chaitanya et al., 2010; Park et al., 2020). Interestingly, poly-ADP ribose levels are increased in the cerebrospinal fluid of PD patients, and some have suggested PARP inhibitors as a potential therapeutic strategy in PD (Kam et al., 2018). Further studies into the underlying mechanism of LRRK2-G2019S leading to increased PARP cleavage in macrophages after membrane damage are of future importance in PD research.

Finally, we were able to recapitulate our findings from mouse primary macrophages in human iPSDM derived from a PD patient carrying the LRRK2-G2019S mutation. Given that we were able to generate isogenic control iPSDM for use as a control in these experiments, our data strongly indicate that these effects on cell death are specific to the LRRK2-G2019S mutation. Future studies exploring the phosphoproteomics of LRRK2 substrates in these iPSDM, and the mode of cell death activated in these cells after LLOMe, would be of interest. We postulate that in resting macrophages the G2019S mutation has a subtle effect and compensatory mechanisms of membrane repair occurring in these basal conditions can compensate. However, when membrane damage is triggered, macrophages undergo cell death due to deficiencies in lysosomal repair. These results could be relevant in the context of macrophage lysosomal function in PD.

## Materials and Methods

### Animals

Depending on the mouse strain, breeding pairs, embryos or sperm were purchased from Taconic Biosciences or Jackson laboratories. The strains utilised were C57BL/6NTac (Tac-WT) and C57BL/6-Lrrk2^tm4.1Arte^ (Tac-G2019S), C57BL/6NJ (NJ-WT), B6(SJL)-Lrrk2^tm4.1Mjff/J^ (NJ-D1994A). Mice were re-derived, bred in-house and maintained under specific pathogen-free conditions at the Francis Crick Institute (UK). Animal studies and breeding were approved by the Francis Crick Institute ethical committee and performed under U.K Home Office project license (P4D8F6075). All animal studies were ethically reviewed and carried out in accordance with Animals (Scientific Procedures) Act 1986.

### Culture of murine primary bone marrow-derived macrophages

Bone marrow was isolated by flushing the femur and tibia of 6-10 weeks female mice with ice-cold PBS as described previously (Schnettger and Gutierrez, 2017). Bone marrow cells were differentiated in RPMI 1640 medium (72400-021, Gibco) containing 10% fetal bovine serum (FBS) (10270-106, Gibco) and 50 ng/ml GM-CSF (576306, BioLegend) for 7 days. For immunofluorescence and live cell snapshot imaging, 6 × 10^4^ cells were seeded per well in 96-well olefin-bottomed plates (6055302, Revvity) and for Western Blotting, 5 × 10^5^ cells were seeded per well in 12-well plates (353043, Falcon) or 1 × 10^6^ cells were seeded per well in 6-well plates (353046, Falcon).

### Gene editing to generate isogenic iPSC

Isogenic control iPSCs were obtained using the CRISPR/Cas9 system. We designed a two-step approach. Briefly, nucleofection using the Amaxa 4D-Nucleofector (V4XP-3024, Lonza) was performed in the LRRK2-G2019S iPSCs (clone ID: CDI00002173) obtained from the Michael J Fox Foundation (MJFF) Parkinson’s Progression Markers Initiative (PPMI) resource. For each nucleofection, 1 × 10^6^ iPSCs were resuspended in 100 μl of P3 buffer (V4XP-3024, Lonza) containing 20 μg Alt-R® S.p. Cas9 Nuclease V3 (1081059, IDT) mixed with 16 μg targeting synthetic chemically modified single guide RNAs (Synthego) (**Table 1**, LRRK2-1step), as (Skarnes et al., 2019). At this point, we screened and selected only the clones containing InDel in the allele harbouring the G2019S mutation by using the ICE platform from Synthego (https://ice.synthego.com/#/). Then, we designed new sgRNA targeting the allele containing the InDel (**Table 1**, LRRK2-2step) and performed nucleofection where 1 × 10^6^ of the selected clone were resuspended in 100 μl of P3 buffer (V4XP-3024, Lonza) containing 20 μg Alt-R® S.p. Cas9 Nuclease V3 (1081059, IDT), mixed with a total of 16 μg of the sgRNAs and 200 pmol of ssODN (**Table 1**), as described before (Skarnes et al., 2019). We screened the clones by Sanger sequencing and selected clones with the wild-type sequence. As a result, we obtained two iPSC clones for experiments: LRRK2-G2019S and isogenic control iPSCs (clone IDs CDI00002173 and 0043052321, respectively). Mycoplasma, Sanger sequencing and Karyostat quality checks were performed in both clones. Additionally, the isogenic clone was tested for pluritest and low pass sequencing. All clones were genotyped and banked for long-term storage.

**Table 1.**
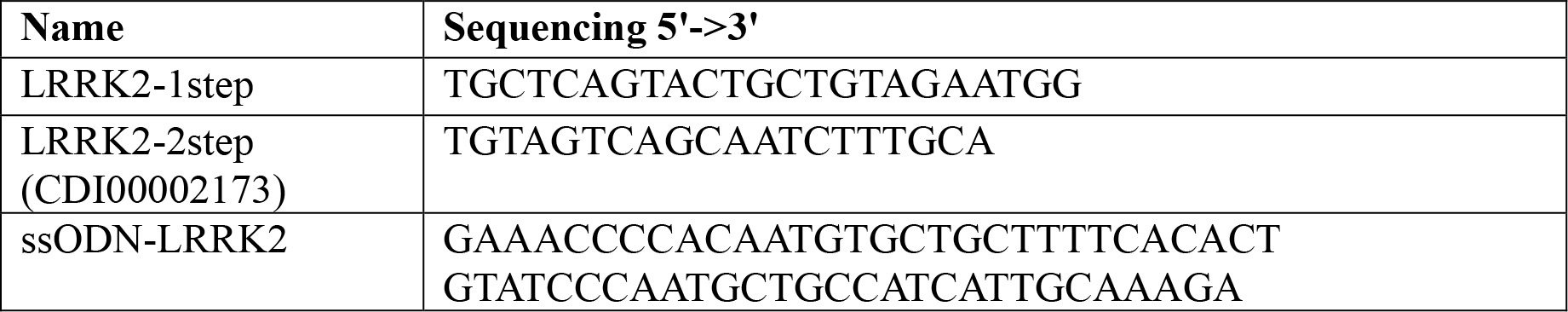
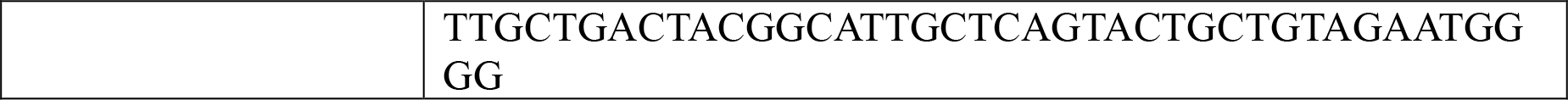
Sequence of sgRNA and ssODN used in the study.

### Culture of induced pluripotent stem cell (iPSC) and iPSC-derived macrophages (iPSDM)

iPSCs were used to generate iPSDM following a previously reported protocol using embryoid bodies (EB) (Bernard et al., 2020; van Wilgenburg et al., 2013). Briefly, EBs were fed daily with two 50% media changes comprising E8 media (A1517001, Life Technologies), BMP4 50 ng/ml (120-05, Peprotech) and VEGF 20 ng/ml (100-20, Peprotech). On day 3, EBs were transferred to T175cm^2^ flasks for macrophage differentiation using factory media consisting of X-VIVO 15 (LZBE02-061Q, Lonza), 2 mM Glutamax (35050061, Life Technologies), 50 μM β-Mercaptoethanol (31350010, Life Technologies), 100 ng/ml M-CSF (300-25, Peprotech) and 25 ng/ml IL-3 (200-03, Peprotech). 20 ml fresh factory media was added once per week and after 4-5 weeks macrophage precursors were collected from the supernatant. These precursors were differentiated into macrophages in X-VIVO 15 (LZBE02-061Q, Lonza) containing 2 mM Glutamax (35050061, Life Technologies) and 100 ng/ml M-CSF (300-25, Peprotech) for 7 days. Macrophages were detached using Versene (15040066, Life Technologies), centrifuged at 300 x g and plated for experiments.

### Antibodies

The following primary antibodies were used for western blot, immunofluorescence, and flow cytometry:

| **Target** | **Host species** | **Company** | **Catalogue number** | **Dilution** | **Application** |
| --- | --- | --- | --- | --- | --- |
| Rab8 | Rabbit | Cell Signaling | 6975 | 1:1000 | Western blot |
| Rab8 pT72 | Rabbit | Abcam | ab230260 | 1:1000 | Western blot |
| Rab10 | Rabbit | Cell Signaling | 8127 | 1:1000 | Western blot |
| Rab10 pT73 | Rabbit | Abcam | ab230261 | 1:1000 | Western blot |
| Rab12 | Rabbit | ProteinTech | 18843 | 1:1000 | Western blot |
| Rab12 pS105 | Rabbit | Abcam | ab256487 | 1:1000 | Western blot |
| Cleaved caspase-3 (Asp175) | Rabbit | Cell Signaling | 9664T | 1:1000 | Western blot |
| Phospho-RIP3 (Thr231/Ser232) | Rabbit | Cell Signaling | 91702 | 1:1000 | Western blot |
| Cleaved Gasdermin D (Asp275) | Rabbit | Cell Signaling | 10137 | 1:1000 | Western blot |
| Cleaved PARP (Asp214) | Rabbit | Cell Signaling | 94885 | 1:1000 | Western blot |
| Phospho-MLKL (Ser 345) | Rabbit | Cell Signaling | 37333S | 1:1000 | Western blot |
| Cleaved-IL-1β (Asp117) | Rabbit | Cell Signaling | 63124S | 1:1000 | Western blot |
| RIP3 | Rabbit | Cell Signaling | 95702S | 1:1000 | Western blot |
| MLKL | Rabbit | Cell Signaling | 37705S | 1:1000 | Western blot |
| LAMP1 | Rat | DSHB | 1D4B | 1:1000 | Western blot |
| β-Actin-HRP | Mouse | Cell Signaling | 12262S | 1:5000 | Western blot |
| CHMP4B | Rabbit | ProteinTech | 13683 | 1:250 | Immunofluorescence |
| Galectin-3 | Rat | BioLegend | 125410 | 1:250 | Immunofluorescence |
| PCD6IP (ALIX) | Rabbit | ProteinTech | 12422 | 1:100 | Immunofluorescence |
| F4/80 | Rat | BioLegend | 123113 | 1:80 | Flow cytometry |
| CD11b | Rat | BioLegend | 101237 | 1:20 | Flow cytometry |
| MHCII | Rat | BioLegend | 107607 | 1:80 | Flow cytometry |
| CD11c | Armenian Hamster | BioLegend | 117319 | 1:200 | Flow cytometry |
| TLR2 | Mouse | BioLegend | 153009 | 1:20 | Flow cytometry |
| CD80 | Armenian Hamster | BioLegend | 104717 | 1:200 | Flow cytometry |
| CD206 | Rat | BioLegend | 141717 | 1:20 | Flow cytometry |
| CD14 | Mouse | BD Biosciences | 562689 | 1:20 | Flow cytometry |
| CD16 | Mouse | BD Biosciences | 557710 | 1:20 | Flow cytometry |
| CD119 | Mouse | BD Biosciences | 558934 | 1:50 | Flow cytometry |
| CD163 | Mouse | BD Biosciences | 563697 | 1:20 | Flow cytometry |
| CD206 | Mouse | BD Biosciences | 561763 | 1:5 | Flow cytometry |
| Siglec-1 (CD169) | Mouse | BD Biosciences | 565248 | 1:20 | Flow cytometry |
| CD86 | Mouse | BD Biosciences | 562433 | 1:20 | Flow cytometry |

For western blot, secondary antibody used was anti-rabbit-HRP conjugate (W410B, Promega) (1:5000 dilution). For immunofluorescence secondary antibodies used were Alexa fluor 488 goat anti-rabbit IgG (H+L) (A11034, Life Technologies) and Cy3 goat anti-rabbit IgG (H+L) (A10520, Life Technologies) (1:800 dilution).

### LLOMe, MLi-2 and silica treatment

LLOMe (4000725, Bachem) was prepared in methanol at a stock concentration of 333 mM and frozen at -20°C. Cells were treated with LLOMe at a concentration of 1 mM by dilution in RPMI (BMDMs) or X-VIVO 15 (iPSDM). 0.3% methanol in RPMI or X-VIVO 15 was used in all control samples. Cells were pre-treated with MLi-2 (5756/10, Tocris) at 100 nM for 2 hours and all subsequent treatments were performed in the presence of MLi-2. Stock crystalline silica (MIN-U-SIL-15, US Silica) was prepared at 30 mg/ml by dilution in PBS immediately prior to use. A working solution of 300 μg/ml crystalline silica was prepared in RPMI for subsequent treatments.

### Phosphoproteomics

For LLOMe treatment, 5 × 10^5^ cells/ml were plated in three 50 mm non-treated tissue-culture dishes (122-17, Thermo Scientific). For control, 6 × 10^5^ cells/ml were plated in a 90 mm petri dish (101R20, Thermo Scientific). Cells were treated with LLOMe for 30 minutes at a concentration of 1 mM by dilution in RPMI medium. Following treatment, cells were washed once in ice-cold PBS and then detached with ice-cold PBS containing Sigma Phosphatase Inhibitor Cocktail 3 (539134, Merck), CoMPLETE Protease Inhibitor tablets (11873580001, Merck), Pierce Phosphatase Inhibitor Mini Tablets (A32957, Thermo Fisher Scientific), 500 nM Okadaic acid (A4540, Stratech) and 1 mM EDTA (11568896, Invitrogen). A pellet was collected by centrifugation (350 xg for 5 minutes) and the pellet was flash-frozen using dry ice/iso-propanol. Four replicates were collected per condition. Pellets were lysed in Urea lysis buffer (8 M urea, 50 mM HEPES pH8.2, 10 mM glycerol-2-phosphate, 50 mM NaF, 5 mM sodium pyrophosphate, 1 mM EDTA, 1 mM EGTA, 1 mM sodium vanadate, 1 mM DTT, 2X cOmplete protease inhibitor cocktail (Merck), 1X phosphatase inhibitor cocktail 3 (Sigma), 500 nM okadaic acid, 1 μM microcystin-LR). After protein concentration was measured using Pierce BCA assay, equal amount (between 165-200 μg) was used of each sample for TMTpro multiplexed quantitative proteomics. Each sample was reduced with 10 mM DTT for 1 h at 56°C and alkylated with 20 mM IAA for 30 min in the dark at room temperature. The reaction was quenched with 20 mM DTT then diluted to <2 M urea with 50 mM HEPES pH8.5. Each resuspended sample was digested with 3.75 μg LysC (Lysyl endopeptidase, 125-05061, FUJIFILM Wako Chemicals) and 12.5 μg Trypsin (MS grade, 90058, ThermoFisher Scientific) at 37°C shaking overnight. Each sample was cleaned-up using Nest Group BioPureSPN MACRO (Proto 300 C18; Part# HMM S18V) and vacuum-dried. Samples were then tandem mass tag (TMT) labelled for an hour at room temperature using a TMTpro 16plex Isobaric Label Reagent Set (0.8 mg per tag, A44520, ThermoFisher Scientific; LOT VE299607) and following manufacturer’s instructions. A small aliquot of each sample was collected for a labelling efficiency and mixing accuracy quality checks (QCs) by liquid chromatography tandem mass spec (LC-MS/MS) using Orbitrap Eclipse Tribrid mass spectrometer and a 60 min HCD MS2 fragmentation method. The rest of the sample was stored at -80°C until QC results. Labelling efficiency of higher than 99% was obtained for each reaction and a mixing accuracy with lower than 1.5x difference between samples with lowest and highest summed intensity. Samples were defrosted, quenched with hydroxylamine for 15 min at room temperature and pooled together. Combined mixture was partially vacuum-dried and acidified to pH 2.0 followed by sample clean-up using C_18_ Sep Pak 1cc Vac, 100 mg bed volume (Waters). Final mixing check was performed by LC-MS/MS with a 240 min HCD MS2 fragmentation method. The peptide mixture was then subjected to high-select sequential enrichment of metal oxide affinity chromatography (SMOAC) to capture phospho-peptides. It was first passed through a high-select TiO_2_ phospho-enrichment column (Thermo Scientific, A32993) following manufacturer protocol. Flow-through and wash fractions were combined, dried, and subsequently used for Fe-NTA phospho-enrichment (Thermo Scientific, A32992). One tenth of the combined flow-through and wash fractions from this enrichment was used for total proteome analysis. The eluates from SMOAC were freeze-dried, solubilised and pooled together. Both total proteome and phospho-proteome samples were subjected to high pH reversed phase fractionation (Thermo Scientific, 84868), dried and resolubilised in 0.1% TFA prior to LC-MS/MS. Total proteome was analysed on Orbitrap Eclipse Tribrid (Thermo) mass spectrometer using 180 min HCD MS2 fragmentation method and 180 min real-time search (RTS) MS3 method (Schweppe et al., 2020). The phosphoproteome was analysed using 180 min HCD MS2 and 180 min MSA SPS MS3 as described in (Jiang et al., 2017). Xcalibur software was used to control the data acquisition. The instrument was run in data dependent acquisition mode.

Raw data were processed using MaxQuant v2.1.3.0 and Uniprot mouse reference proteome from March 2021. Processed data were then analysed using an R-coding script tailored to isobaric labelling mass spectrometry. The script was generated as a hybrid using the backbone and differential gene expression analysis of ProteoViz package (Storey et al., 2020) as a general script workflow and borrowing the normalization script from Proteus package (Marek et al.). Briefly, the “proteinGroups.txt” and “Phospho (STY)Sites.txt” tables were read into matrices and filtered for “reverse” hits, “potential contaminant” and proteins “only identified by site”. TMT LOT-corrected reporter intensities were then normalized using CONSTANd (Van Houtven et al., 2021), log2-transformed and differentially analysed using Linear Models for Microarray Data (limma). Volcano plots were generated using ggrepel (a ggplot2 extension) as part of the tidyverse.

### SDS-PAGE and western blot

Cells were washed once with PBS and lysed on ice in RIPA buffer (Millipore, 20-188) containing protease and phosphatase inhibitor (Thermo Scientific, 78440). Protein levels were normalised using the Pierce BCA protein assay kit (23227, Thermo Scientific) as per manufacturer’s instructions. Samples were boiled at 70°C for 15 minutes in LDS sample buffer (Thermo Fisher Scientific, NP008) and NuPage Sample Reducing Agent (Thermo Fisher Scientific, NP009). Samples were loaded into 4–12% Bis-Tris gel (Thermo Fisher Scientific, WG1403), and electrophoresis was performed at 100 V for 120 minutes. The gels were transferred onto a PVDF membrane using an iBlot2 (Thermo Fisher Scientific, IB21001) using program P0. Membranes were blocked in 5% skimmed milk powder in TBS plus 0.05% Tween20 (TBS-T) for 1 hour at room temperature, then incubated with primary antibody overnight at 4 °C. For detection of phosphorylated proteins, membranes were blocked in 5% BSA (Cell Signaling, 9998S) in TBS-T. Membranes were washed in TBS-T and incubated with horseradish peroxidase (HRP)-conjugated secondary antibodies for 1 hour at room temperature. Membranes were developed with enhanced chemiluminescence reagent (Bio-Rad) and imaged on an Amersham GE Imager 680 (GE Healthcare). The molecular weight ladder was from Abcam (116028). Western blots were quantified by densitometry using FIJI software.

### Indirect immunofluorescence

BMDMs were seeded in 96 well plates (6055302, Revvity) and then treated with vehicle or LLOMe 1 mM. After 30 minutes, cells were fixed in 4% PFA (15710, Electron Microscopy Sciences) for 15 minutes and then washed once with PBS. The samples were quenched with 50 mM NH4Cl (A9434, Sigma) for 10 minutes at room temperature and permeabilised in ice-cold methanol for 10 minutes at -20°C. Primary antibodies were diluted in PBS containing 5% FBS (10270-106, Gibco) and incubated for 1 hour at room temperature. Samples were washed three times in PBS and then secondary antibody was added for 1 hour at room temperature. Following three washes with PBS, nuclear staining was performed using 300 nM DAPI (D3571, Life Technologies) in PBS for 10 minutes. After a further three washes in PBS, the samples were imaged using an automated confocal high-content imaging system (Revvity, Opera Phenix High-Content Screening System) with 63x 1.15 NA water-immersion objective and excitation lasers (405 nm, 488 nm, 561 nm, 640 nm) and preset emission filters. Channels were separated during acquisition to prevent fluorescence crosstalk. At least 800 cells per condition were imaged.

Image analysis was performed using Harmony 4.9 image analysis software (Revvity). First, stacks were processed to create a maximum projection image. Nuclei were segmented using the “Find nuclei” building block (method B). Following this, cell boundaries were segmented using the “Find Cytoplasm” building block (method A). Cell area was then calculated using the “Calculate Morphology Properties” building block. Galectin-3, CHMP4B and Alix puncta were segmented using the “Find Spots” building block (method D) followed by the “Select Population” building block (puncta filtered on basis of “corrected spot intensity”). Total puncta area was calculated using the “Calculate Morphology Properties” building block. Finally, the total puncta area per cell was calculated using the “Calculate Properties” building block.

### LysoTracker leakage assay

BMDMs were seeded in 96 well plates (6055302, Revvity) and stained with 25 nM LysoTracker DND-99 (LTR) (L7528, Thermo Fisher Scientific) for 30 minutes. Following this, live cell snapshot imaging was performed at 37°C, 5% CO2 using an automated confocal high-content imaging system (Revvity, Opera Phenix High-Content Screening System) with 40x 1.1 NA water-immersion objective, 561 nm excitation laser and 570-630 nm emission filter at a power of 30% and 100ms exposure time. Cells were imaged once prior to any treatment to establish a baseline. Then, 1 mM LLOMe in RPMI was added and the cells were imaged every minute for 5 minutes to track lysosomal damage. Following this, cells were washed twice in RPMI and fresh media containing 25 nM LTR was added, and the cells were imaged every minute for 10 minutes to track lysosomal recovery. All media changes and wash steps were performed without moving the plate so that the same cells were imaged at each snapshot. Image analysis was performed using Harmony 4.9 image analysis software (Revvity). First, stacks were processed to create a maximum projection image. Cells were segmented using the “Find Cells” building block (method P). LTR puncta were identified using the “Find Spots” building block (method A) and LTR spot area and intensity were calculated using the “Calculate Morphology Properties” and “Calculate Intensity Properties” building blocks. The mean LTR cytoplasmic intensity at each time-point was normalised to the baseline LTR cytoplasmic intensity to correct for differences in baseline prior to analysis.

### High-content live imaging of BMDM and iPSDM cell death

Cells were seeded in 96 well plates (6055302, Revvity) and stained with Blue/Green ReadyProbes Cell Viability Imaging Kit (Invitrogen, R37609) according to manufacturer’s instructions. Cells were treated with LLOMe 1 mM, silica 300 μg/ml or vehicle just prior to the start of imaging. Live cell snapshot imaging was performed at 37°C, 5% CO2 using an automated confocal high-content imaging system (Revvity, Opera Phenix High-Content Screening System) with 40x 1.1 NA water-immersion objective, and excitation lasers (405 nm and 488nm) and preset emission filters. The 405 nm laser was used at 20% power with 100 ms exposure. The 488nm laser was used at 5% power with 80 ms exposure. The channels were separated during acquisition to prevent fluorescence crosstalk. Snapshots were taken every 15 or 30 minutes depending on the length of acquisition and treatment. Image analysis was performed using Harmony 4.9 image analysis software (Revvity). First, stacks were processed to create a maximum projection image. The total cell number was calculated using the “Find nuclei” building block in the NucBlue channel (method B). Then, intensity of NucGreen within each nucleus was calculated using the “Calculate Intensity Properties” building block. The total number of dead cells was calculated by setting a threshold for NucGreen intensity and segmenting this population using the “Select Population” building block. The percentage of dead cells was calculated by dividing the total number of NucGreen+ nuclei over the total number of NucBlue nuclei.

### Induction of cell death controls and LPS treatment

Where indicated, LPS 250 ng/ml (L8274, Sigma) was added for 3 hours prior to downstream applications. For induction of apoptosis, BMDMs were stimulated with 100 ng/ml TNF-α (300-O1A, Peprotech) and 125 nM 5Z-7-Oxozeaenol (9890, Sigma) for 7 hours. For induction of necroptosis, BMDMs were stimulated with 100 ng/ml TNF-α (300-O1A, Peprotech), 125 nM 5Z-7-Oxozeaenol (9890, Sigma) and 10 μM Z-VAD-FMK (tlrl-vad, Invivogen) for 7 hours. For induction of pyroptosis, BMDMs were pre-treated with 250 ng/ml LPS (L8274, Sigma) for 3 hours followed by 20 μM Nigericin for 3 hours (ICT9146, Bio-Rad).

### Flow cytometry

1 × 10^6^ cells were pelleted by centrifugation (350 x g for 5 minutes at 4°C), washed with cell staining buffer (420201, BioLegend) and incubated with TruStain fcX™ (156604, BioLegend) for 20 minutes at 4°C. Cells were resuspended in cell staining buffer with fluorochrome-conjugated antibodies for 30 minutes at room temperature in the dark. Cells were washed in cell staining buffer and resuspended in 500 μL cell staining buffer for acquisition. Data were acquired using an BD LSRFortessa™ Cell Analyzer (BD Biosciences) and analysed using FlowJo software (FlowJo, LLC, v10.8.0). At least 10,000 events per condition were recorded.

### Bafilomycin treatment and DQ-BSA assay

Cells were seeded in 96 well plates (6055302, Revvity) and incubated with 100 nM Bafilomycin A1 (Merck, B1793) or vehicle (DMSO) for 2 hours. Next, 10 μg/ml DQ-BSA (Thermo Fisher Scientific, D12051) was added for 4h. The cells were then washed x3 in RPMI (BMDM) or X-VIVO 15 (iPSDM), counterstained with NucBlue and imaged at 37°C, 5% CO2 using an automated confocal high-content imaging system (Revvity, Opera Phenix High-Content Screening System) with 40x 1.1 NA water-immersion objective, and excitation lasers (405 nm and 561 nm) and preset emission filters. The 405 nm laser was used at 30% power with 100 ms exposure. The 561 nm laser was used at 20% power with 200 ms exposure. Image analysis was performed using Harmony 4.9 image analysis software (Revvity). First, stacks were processed to create a maximum projection image. Cells were segmented using the “Find Nuclei” (method B) and “Find Cytoplasm” (method A) building blocks. Following this, intensity of DQ-BSA in the cytoplasm was calculated using the “Calculate Intensity Properties” building block.

### Statistical analysis

Statistical analyses were performed using GraphPad Prism software version 9.3.1. Data are shown as mean ± SD. Unpaired t-tests were performed for statistical analysis of experiments with two groups. For more than two groups, analysis was performed using one-way analysis of variance (ANOVA) with Šidák’s multiple comparisons test or two-way ANOVA with Šidák’s multiple comparisons test, as indicated in the figure legends.

## Acknowledgments

This work was supported by the Francis Crick Institute (to MGG), which receives its core funding from Cancer Research UK (FC001092), the UK Medical Research Council (FC001092), and the Wellcome Trust (FC001092). This project has received funding from the Michael J Fox Foundation and the and The Parkinson’s Progression Markers Initiative.

## Author contributions

MGG conceived the project. HM helped at initial stages of the project. RM performed most of the experiments and analysed the data. RM, SM, CHL, AR, NA, EP, SU performed experiments, generated tools, developed the methods and analysed the data. MGG and RM wrote the manuscript and prepared the figures. All the authors provided feedback on the manuscript.

