## Supplementary figures for "LRRK2 kinase dependent and independent function on endolysosomal repair promotes macrophage cell death"

**Supplementary Figure 1**

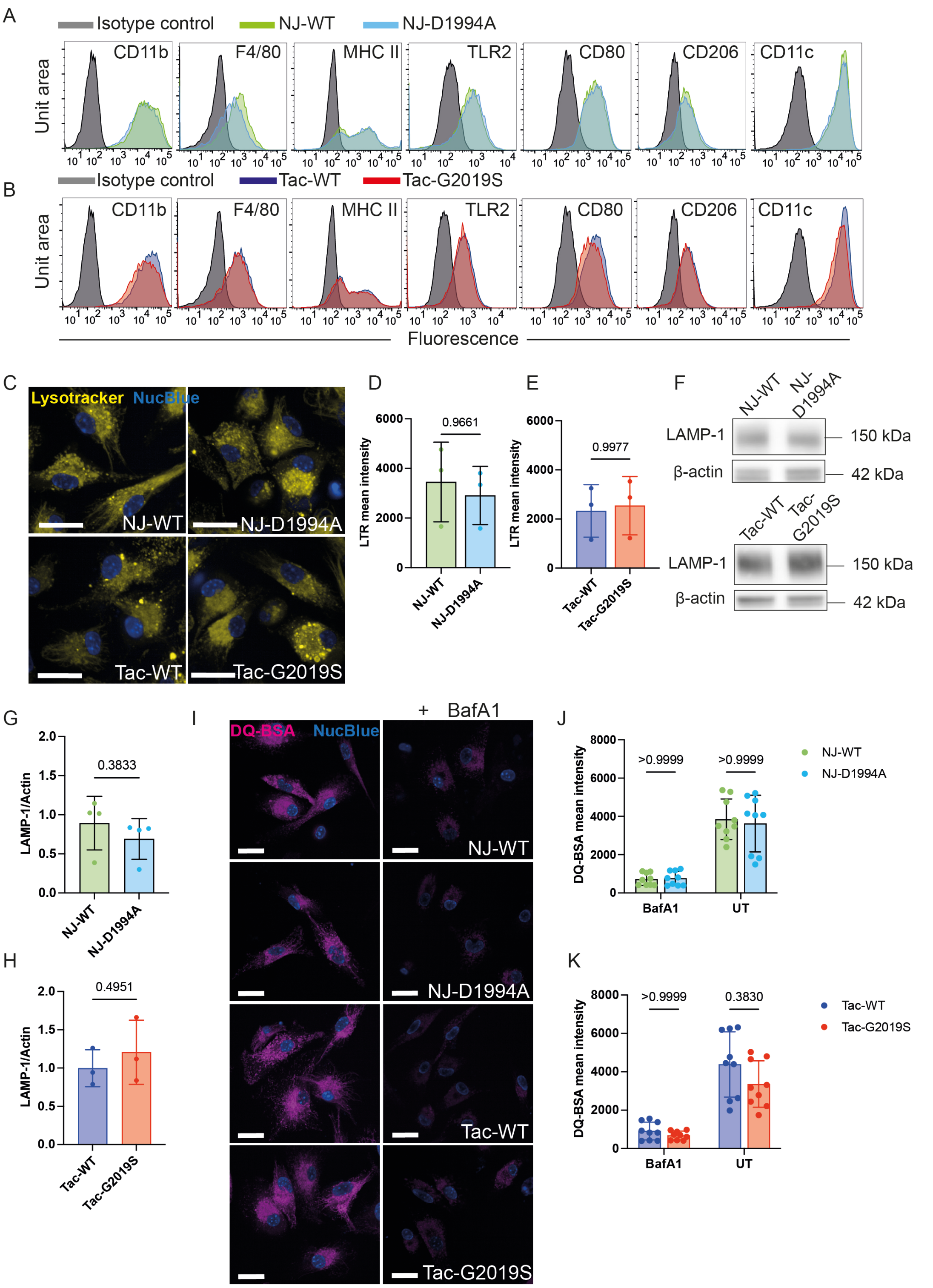


**Supplementary Figure 1. Immunophenotyping and characterisation of lysosomal activity from LRRK2-associated genotypes.** (A and B) Flow cytometry characterisation of surface expression of macrophage markers in BMDM. Data is from one representative experiment. Data is presented as histograms with compensated fluorescence of the indicated surface marker on the x-axis. (C) Representative images of live BMDM stained with Lysotracker (LTR) and NucBlue dye. Scale bars = 20 μm. (D and E) Quantitative analysis of the mean LTR cytoplasmic intensity. Data shown is mean ± SD from 3 independent experiments (n=5 independent wells). One-way ANOVA followed by Šidák's multiple comparisons test. (F) Western blot analysis of LAMP-1 and β-actin. (G and H) LAMP-1 band intensity weas quantified by densitometry and normalised to β-actin. Data are mean ± SD from 4 independent experiments. Two-tailed t-test. (I) Representative images of live BMDM pre-treated with Bafilomycin A1 (BafA1) 100 nM followed by incubation with DQ-BSA and NucBlue dye. Scale bars = 20 μm. (J and K) Quantitative analysis of the mean DQ-BSA cytoplasmic intensity. Data shown is mean ± SD from 3 independent experiments (n=9 independent wells). Two-way ANOVA followed by Šidák's multiple comparisons test.

### Supplementary Figure 2

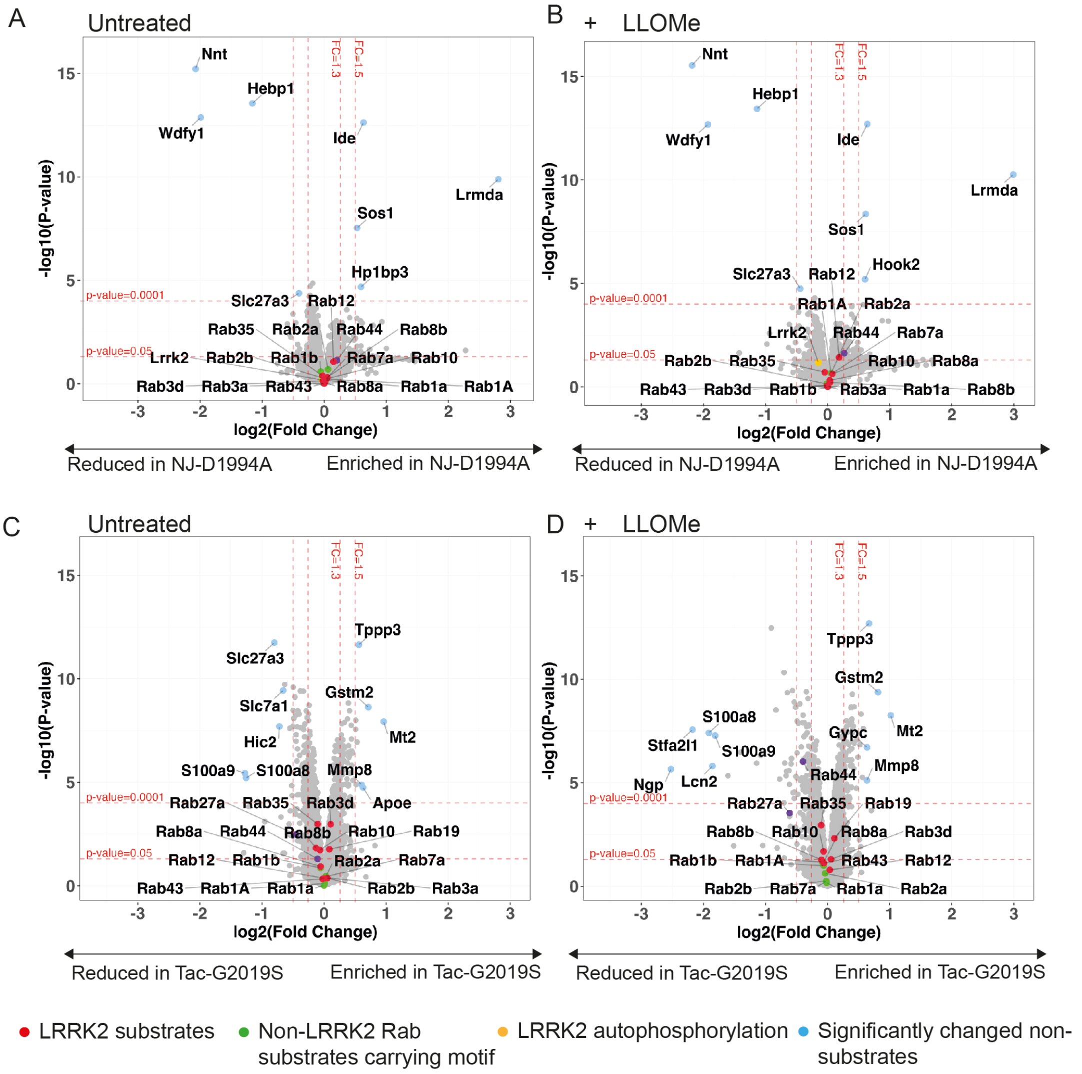


**Supplementary Figure 2. Total proteome in untreated and LLOMe-treated conditions from LRRK2 associated genotypes.** (A to D) BMDMs were treated with LLOMe 1 mM for 30 minutes and cells were analysed by mass spectrometry. Volcano plots in the untreated (A and C) and LLOMe-treated (B and D) conditions of the difference in total protein levels between (A to B) NJ-WT and NJ-D1994A BMDMs and (C to D) Tac-WT and Tac-G2019S BMDMs. Each point represents one protein. The x-axis shows the $log_2$-transformed fold change and the y-axis shows the significance by $-log_{10}$-transformed P-value, obtained by linear models for microarray data. A fold-change greater than 1.3 and p value<0.05 was deemed significant.

### Supplementary Figure 3

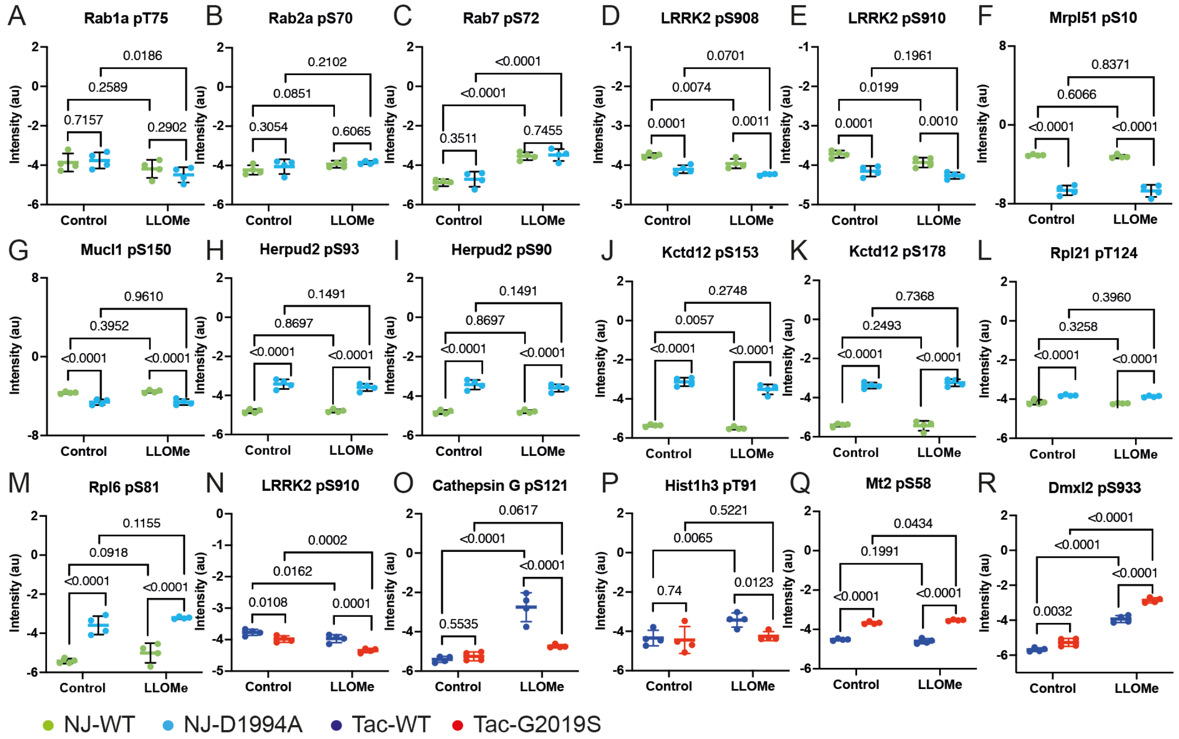


**Supplementary Figure 3. Phosphoproteomic analysis of non-LRRK2 Rab substrates carrying motif, LRRK2 phosphorylation sites and significantly altered non-substrates.** Scatterplots showing the raw intensity data obtained by mass spectrometry and p-values for: non-LRRK2 Rab substrates carrying the switch II motif (A to C); LRRK2 phosphorylation sites pS908 and pS910 (D and E, N); significantly altered non-substrates (F to M, O to R). Each point represents the normalised and log2 transformed TMT-corrected reporter intensity value obtained by mass spectrometry (n=4 biological replicates). P values by linear models for microarray data.

### Supplementary Figure 3

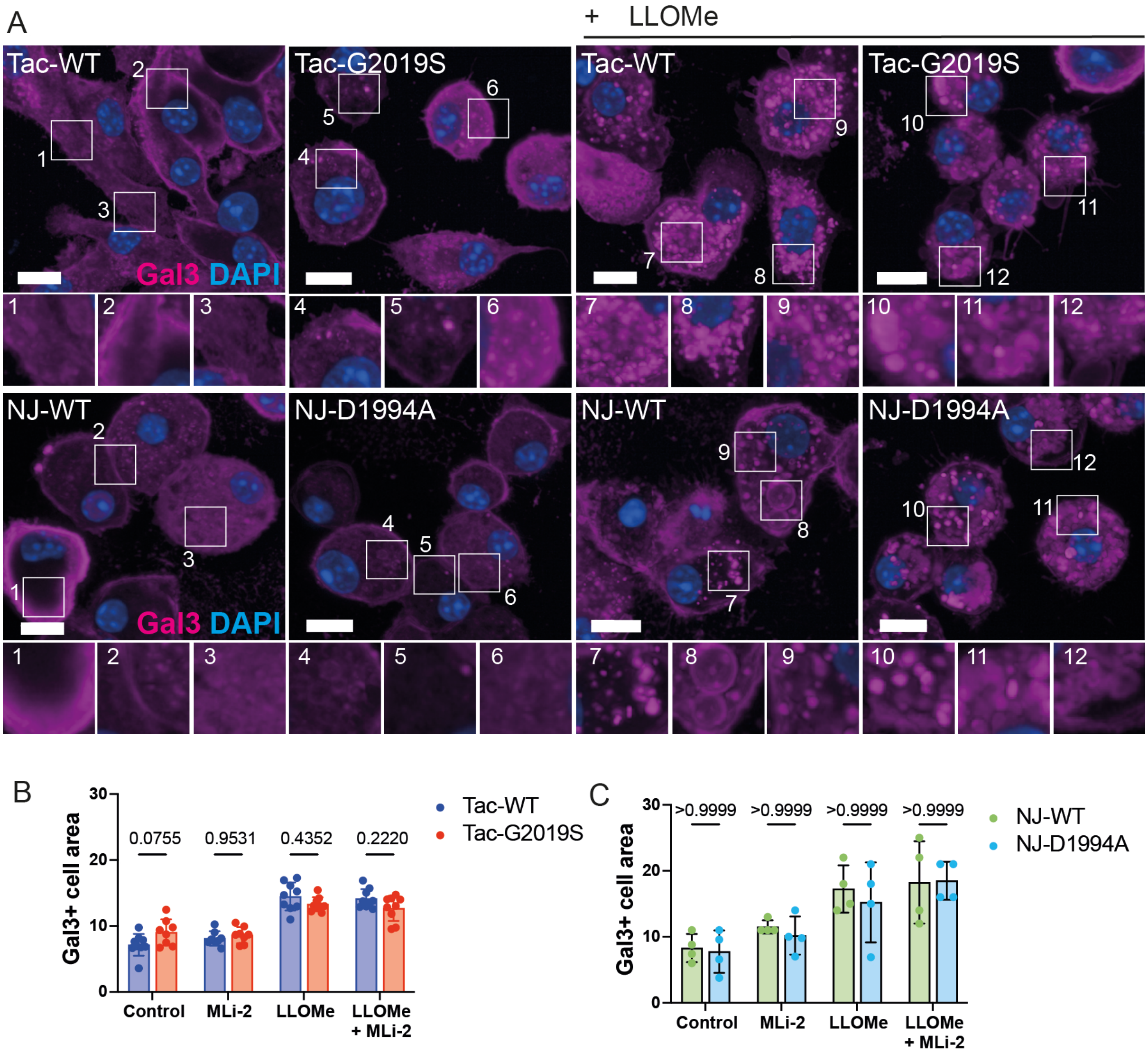


**Supplementary Figure 4. Lysosomal damage after LLOMe is unchanged by LRRK2 kinase activity.** (A) BMDMs were treated with LLOMe 1 mM for 30 minutes and endogenous Galectin-3 (Gal3) positive cell area was visualised by immunofluorescence and quantified using Harmony software. Scale bars, 10 µm. Data representative of (B) 3 independent experiments (n=9 independent wells) and (C) 4 independent experiments. Results are shown as mean ± SD. Two-way ANOVA followed by Šidák's multiple comparisons test.

### Supplementary Figure 5

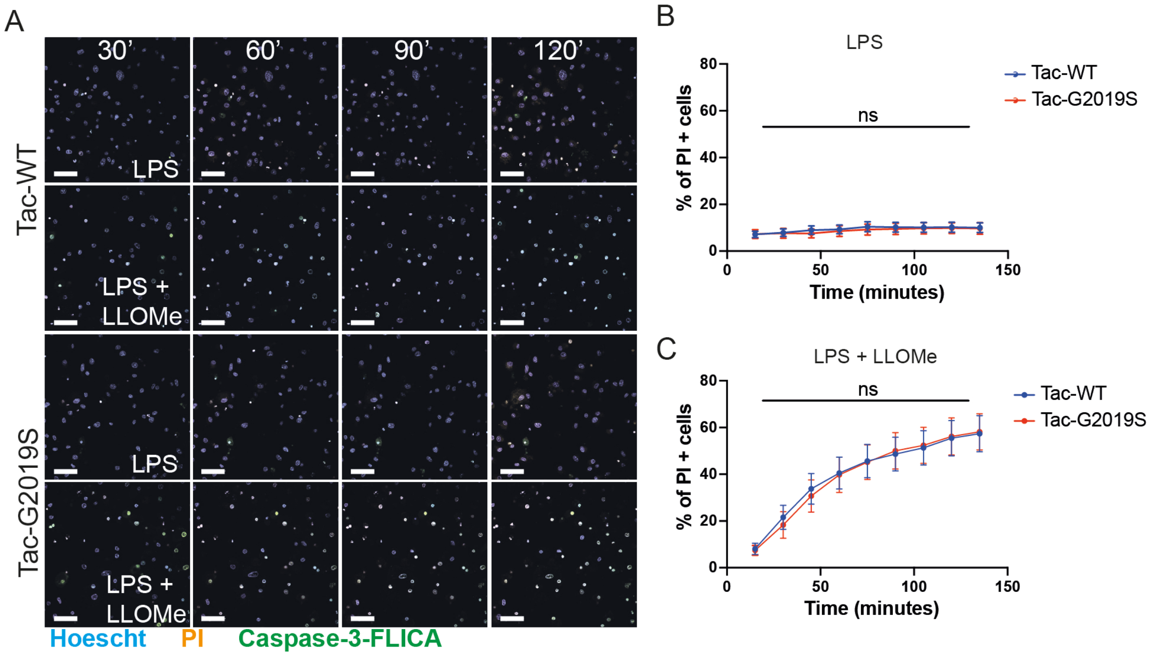


**Supplementary Figure 5. Inflammasome activation by LPS pre-treatment results in similar levels of cell death between WT and LRRK2-G2019S after lysosomal damage.** (A) Snapshot of live BMDM pre-treated with LPS 250 ng/ml followed by LLOMe 1 mM and imaged every 30 minutes. Nuclear staining for live/PI/caspase-3 (blue/orange/green). Scale bars = 50 µm. (G to I) Quantitative analysis of the percentage of dead cells (PI-positive) at each timepoint. Data shown is the mean ± SD from three independent experiments. Two-way ANOVA followed by Šidák's multiple comparisons test. ns = non-significant.

### Supplementary Figure 6

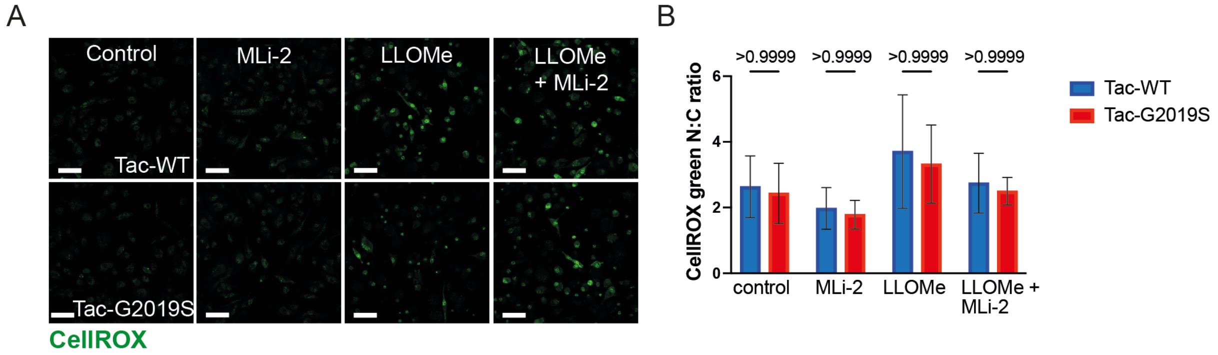


**Supplementary Figure 6. Cellular reactive oxygen species are unchanged after lysosomal damage in LRRK2-G2019S macrophages.** (A) Live BMDM were pre-treated with MLi-2 100 nM for 2 hours followed by LLOMe 1 mM for 30 minutes. Cells were stained for reactive oxygen species using the CellROX green dye. Scale bars = 50 µm. (E) Quantitative analysis of the mean CellROX green nuclear:cytoplasmic ratio. Data shown is the mean + SD from 3 independent experiments. Two-way ANOVA followed by Šidák's multiple comparisons test.
